# A Hard Sphere Model for Single File Water Transport Across Biological Membranes

**DOI:** 10.1101/2023.10.11.561951

**Authors:** Gerald S. Manning

**Affiliations:** Department of Chemistry and Chemical Biology, Rutgers University, 610 Taylor Road, Piscataway, NJ 08854-8087, USA

## Abstract

We use Gürsey’s statistical mechanics of a one-dimensional fluid to find a formula for the *P*_*f*_ */P*_*d*_ ratio in the transport of hard spheres across a membrane through a narrow channel that can accommodate only single file movement. *P*_*f*_ is the membrane permeability for osmotic flow, and *P*_*d*_ the permeability for exchange across the membrane in the absence of osmotic flow. The deviation of the ratio from unity indicates the degree of cooperative transport relative to ordinary diffusion of independent isolated molecules. In contrast to an early idea that *P*_*f*_ */P*_*d*_ must be equal to the number of molecules in the channel, regardless of the physical nature of the interactions among the molecules, we find a functional dependence on the fractional occupancy of the length of the channel by the hard spheres. We also attempt a random walk calculation for *P*_*d*_ individually, which gives a result for *P*_*f*_ as well when combined with the ratio.

## 1 Introduction

Water transport across membranes can be characterized by two distinct membrane permeabilities, the osmotic permeability *P*_*f*_, and the diffusion, or exchange, permeability *P*_*d*_ [1]. The *P*_*f*_ */P*_*d*_ ratio provides useful information about the mechanism of flow. If it equals unity, as for lipid bilayers, one may assume that water crosses the membrane as widely separated individual molecules. If it is much larger than unity, then water presumably exists inside the membrane in its usual bulk liquid form, and its flow under applied pressure, or, equivalently, in osmosis, is predominantly convective.

Biological membranes are spanned by protein channels, so narrow that water molecules traverse them in single file, molecule after molecule. It might have been expected that the osmotic and diffusion permeabilities would be equal for these channels, but as it turned out, the *P*_*f*_ */P*_*d*_ ratio has been reliably measured as significantly greater than unity, although not as large as for bulk flow through most synthetic membranes. An early idea was that the single-file ratio *P*_*f*_ */P*_*d*_ should equal *N*, the number of water molecules in the column, as following from the mutual impenetrability of the water molecules [1]. However, a focus on the permeability ratio was all but lost before resolution of this question. In this paper we show that the inability of the water molecules in a column to pass each other does not necessarily lead to the answer *N* for the permeability ratio. Nonetheless, we find that the ratio exceeds unity for a hard sphere model (impenetrable balls, all with the same diameter, with no forces between balls when not in contact). This paper supersedes an earlier one by the author that was based on an unrealistic boundary condition [2].

## 2 Background

The osmotic, or filtration, permeability coefficient *P*_*f*_ for the molar water flux *J*_*w*_ across a membrane (number of moles per unit membrane area per second) as induced either by an applied pressure difference Δ*P* or an osmolyte concentration difference Δ*C*_*s*_ (osmolyte concentration is the total concentration of impermeable solute) is defined by the equation,^1^

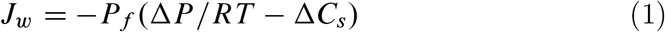

which can be called the Fundamental Law of Osmosis [3]. As follows from Debye’s analysis of the van’t Hoff equation [3,4], an osmolarity difference induces a pressure gradient across the membrane identical [5,6] to the gradient induced by an applied pressure difference [7].

It is of interest to compare the value of *P*_*f*_ with the value of a different permeability coefficient *P*_*d*_ for the same membrane, defined by,

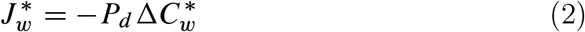

where 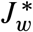 is the molar flux of a tracer water isotope across the membrane as induced by a concentration difference 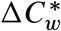 of tracer water with equal pressures on both sides of the membrane and no osmolyte present [1]. From its definition we see that *P*_*d*_ measures the rate of water exchange across the membrane in the absence of any driving forces, and due only to the self-diffusion of the water molecules, that is, their thermal random movement. The conventional name for *P*_*d*_ is the diffusion permeability, and we will stick with convention, although a better nomenclature would perhaps have been the exchange permeability.

For membranes allowing bulk, or convective, flow of water,

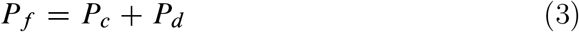

where the total observed osmotic flow is understood to consist of independent convective and diffusive components, the latter due to the stochastic movements of water molecules that are random in the absence of directed forces but biased in the direction of a force such as a pressure gradient [3]. The osmotic permeability *P*_*f*_ can be measured experimentally from its definition, Eq. (1), and also the diffusion permeability from its definition, Eq. (2), but the convective component *P*_*c*_ can only be inferred by difference from Eq. (3). In this way Mauro [8], and Robbins and Mauro [9], were able to find that the convective component strongly dominates in a series of synthetic collodion membranes, indicating the presence of bulk water inside the membrane. The diffusive component is always present, however, and becomes increasingly significant with increasing density of membrane polymer material [9]. For lipid bilayers, the same experimental procedures (once the difficulty of stirring the solution right up to the membrane is overcome [1]) show the absence of a convective component. Both *P*_*f*_ and *P*_*d*_ are independently measured and found to be equal [1], meeting the expectation that water crosses the lipid bilayer as widely separated molecules.

It is useful to rearrange Eq. (3) to the form,

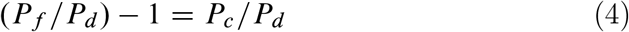

For the synthetic collodion membranes, since measurements show that *P*_*f*_ */P*_*d*_ *>>* 1, it follows that *P*_*c*_ *>> P*_*d*_, in words, convection dominates diffusion. But if *P*_*f*_ */P*_*d*_ = 1, as in lipid bilayers, *P*_*c*_ = 0, and there is no convective water flow.

In the setting of biological cell membranes, a significant part of water exchange and flow is directly across the lipid bilayer, 10–20% [1, 10]. But biological membranes are punctuated by proteins like the aquaporins that provide narrow channels allowing efficient single file flow of water molecules, and these pores carry most of the transmembrane flow. In fact, it was the measurements of *P*_*f*_ */P*_*d*_ ratios, and their observed values substantially greater than unity, that first suggested the existence of these protein channels.

Measured values of the *P*_*f*_ */P*_*d*_ ratio for single file channels are greater than unity, but we would not wish to call this molecule-by-molecule flow predominantly convective, or bulk, as in the case of most synthetic membranes containing bulk liquid water. Can values of *P*_*f*_ *=P*_*d*_ exceeding unity be explained in this case, invoking only ordinary thermal and pressure driven movements of independent molecules, or are specific properties of water molecules necessarily involved, such as their propensity (in an inhospitable carbon nanotube environment, for example [11]) to link by direct hydrogen bonding into a connected chain, like monomers in a polymer? In this paper we explore some dynamics of a single file of independent hard spheres, and find that their flow is consistent with *P*_*f*_ */P*_*d*_ ratios greater than unity. Early efforts in this direction will be reviewed in the Discussion section.

## 3 The *P*_*f*_ */P*_*d*_ ratio in single file flow

We start with the equation,

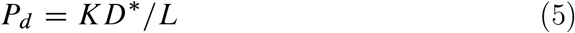

where *D\** is the self-diffusion coefficient of water molecules inside a membrane of thickness *L*. If the water molecules cross the membrane through a protein channel, we assume for simplicity that the length of the channel also equals *L*. With reference to the definition of *P*_*d*_ in Eq. (2), the driving force for tracer diffusion across the membrane on the molecular level is not the concentration difference between tracer concentrations in the reservoirs bathing the membrane but the tracer concentration gradient inside the membrane. Since the factor 1*/L* does not appear explicitly in the definition, it must be included as a factor in any expression for *P*_*d*_. The constant factor *K* accounts for the dependence of the permeability on the amount of water in the membrane relative to the outside reservoirs. The explicit expression for *K*, given subsequently, is not needed in this section.

Next we consider osmotic water flow across the membrane. As discussed above in the Background section, it is caused by a pressure gradient. Thus for a membrane containing liquid water, the flow is predominately bulk, or convective, flow. A lipid bilayer contains a slight amount of water dissolved in it as discrete independent molecules, and in osmosis these molecules are transported across the bilayer by a pressure gradient [3]. The molecules drift in a stochastic diffusive manner in the direction of lower pressure. In single file also, the movement of water molecules in osmosis is best described in terms of the diffusive movement of the discrete molecules in the column, since the water does not exist there in bulk. We therefore postulate the same starting point for *P*_*f*_ as for *P*_*d*_,

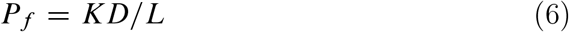

but for *P*_*f*_ the diffusion coefficient *D* may not be equal to the self-diffusion coefficient *D\**. The *P*_*f*_ */P*_*d*_ ratio is therefore,

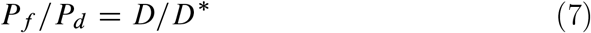

The concentration dependence of diffusion coefficients [12-15] is crucial for present purposes. In self-diffusion, the environment of a thermally agitated molecule is on average symmetric, and the influence of surrounding molecules is not biased in any particular direction. But in diffusive flow caused by a concentration gradient, the more numerous molecules on one side of any given diffusing molecule exert a stronger concentration effect (from intermolecular forces) than the less numerous ones on the other side. Because of this asymmetry, we can write,

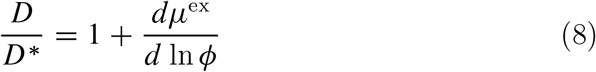

where in this equation *μ*^ex^, in dimensionless units of *k*_*B*_*T*, is the “excess” part of the chemical potential of the diffusing species, which accounts for non-ideal concentration effects, and therefore tends to zero when the concentration is low and intermolecular interactions become negligible. For the concentration derivative in the single-file case of diffusing hard spheres, we have defined *ϕ* as a measure of the linear density of the spheres,

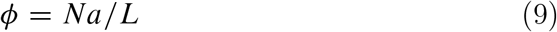

In words, then, *ϕ* is the average fraction of the length of the column occupied by the spheres, each sphere possessing diameter *a*.

The interaction between hard spheres is entirely repulsive (the potential energy of interaction jumps from zero to infinity when two spheres make contact). It follows that the more numerous hard spheres at the high concentration side of a given sphere will push that sphere towards the low concentration side, augmenting self-diffusion. The derivative in Eq. (8) is thus expected to be positive. We confirm the expectation by making an exact calculation. From the statistical mechanical analysis of a linear assembly of molecules as presented in 1950 by Gürsey [16,17], we find for the special case of hard spheres in single file [15,16],

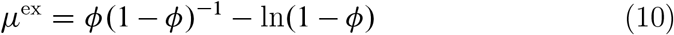

Both terms in this expression increase when *ϕ* increases. When *ϕ* tends to zero, so does *μ*,^ex^.

The result for the *P*_*f*_ */P*_*d*_ ratio follows,

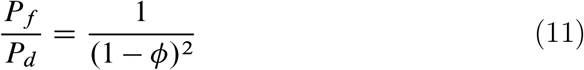

Notice that in this section we did not attempt to calculate *P*_*f*_ and *P*_*d*_ individually but only their ratio, which is plotted in Figure 1. At low densities, a regime where the molecules do not interact at all, so that the effect of asymmetric interactions in osmotic flow vanishes, *ϕ* → 0, and we see that *P*_*f*_ becomes equal to *P*_*d*_. But as *ϕ* increases, the expression for the *P*_*f*_ */P*_*d*_ ratio becomes significantly greater than unity, until finally diverging at *ϕ =* 1, when the spheres are in contact (closest packed) at infinitely high interaction energy, and the slightest asymmetry of a hard sphere environment has a dominantly strong effect.

**Figure 1.**
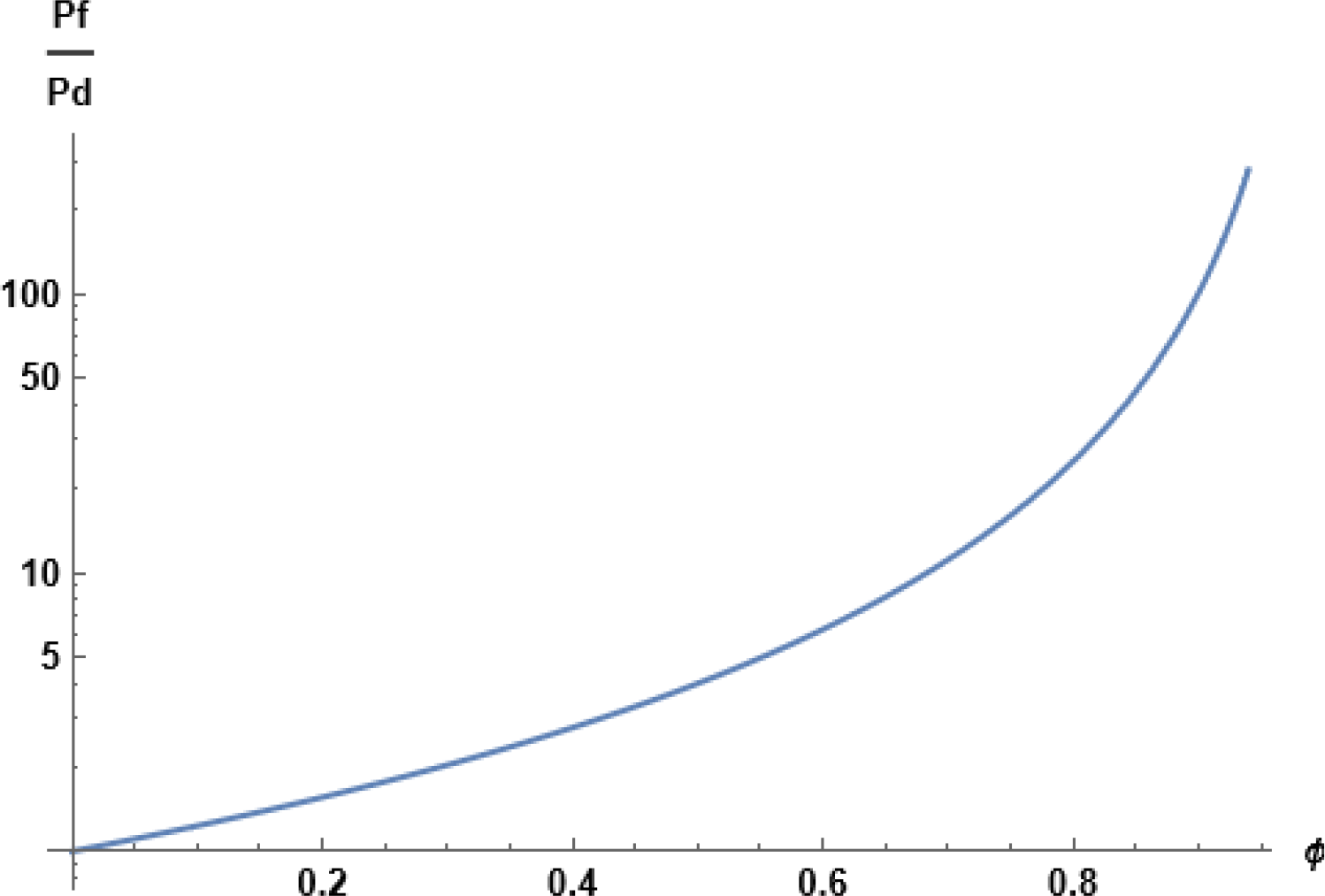
Graphical representation of Eq. 11 for the Pf/Pd ratio as a function of fractional occupancy of single file channel with hard spheres.

Despite the divergence at closest packing, Eq. (11) yields realistic values for the *P*_*f*_ */P*_*d*_ ratio when the density parameter *ϕ* is given in a realistic range. For gramicidin A pores [1], the measured value of *P*_*f*_ */P*_*d*_ is 5.3, which matches Eq. (11) for *ϕ*=0.57. The value of *P*_*f*_ */P*_*d*_ measured [10] for aquaporin is 12.7, matched by Eq. (11) when *ϕ*=0.72. It is worth observing that for values of *ϕ* greater than 0.5, the average single file spacing of hard spheres cannot accommodate another sphere. In other words, in a model such as we have been exploring, it is not necessary for meaningful interpretation of “full occupancy” that *ϕ* be assigned the value unity.

As a further example, we may recall Robbins and Mauro’s measurements of *P*_*f*_ */P*_*d*_ for synthetic collodion membranes [9]. For the membrane most dense in polymer material, the measured value of *P*_*f*_ */P*_*d*_ was 36. The passage of water across membranes of higher density could not be detected. We therefore take the number 36 as a speculative measure of a maximum value that natural selection could have reached for *P*_*f*_ */P*_*d*_ by means of a narrow protein pore in exploiting the advantages of concerted flow over diffusion of isolated molecules into and out of a cell. The corresponding value of *ϕ* from Eq. (11) is 0.83.

Another highly speculative conjecture is that if there were a gating mechanism allowing a cell membrane to regulate the value of the fractional occupancy *ϕ*, then osmotic flow through the membrane could be brought to low levels approaching the rate of water exchange, *P*_*f*_ = *P*_*d*_, simply by allowing *ϕ* to become small. But if *ϕ* had a larger value nearer to unity, then the rate of osmotic flow could be substantially larger than exchange.

## 4 The individual permeabilities

We regard the result for the *P*_*f*_ */P*_*d*_ ratio as our central one. However, at the cost of further modeling, we will also try to find individual expressions for the osmotic and diffusion permeabilities. Actually, we need only an expression for the latter, since *P*_*f*_ will follow immediately from their ratio.

For *P*_*d*_ in Eq. (5), we need expressions for both *K* and *D\**. The water partition coefficient 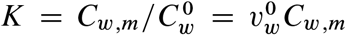, that is, the ratio of water concentrations (number of molecules per unit volume) inside the membrane and in the ideally dilute reservoirs, where 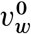 is the volume per molecule of pure water. For the concentration *C*_*w;m*_ of water in the membrane, the volume of the membrane is *AL*, where *A* is the area of a cross section, and if the water crosses through *n* single file channels, each containing *N* hard-sphere “water” molecules, then *C*_*w;m*_ = *nN/AL* = *nϕ=aA*, where we have used the fractional occupancy *ϕ* of the channel, Eq. (9). We therefore have 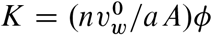.

As in traditional diffusion theory, we describe the thermal movement of the molecules in the channel by a random walk, in this case one-dimensional. The mean free path *l* = (*1 ϕ*)*l*_*0*_, where *l*_*0*_ is the mean free path when *ϕ* →0. In words, *l*_*0*_ is the mean free path for a molecule in a channel so sparsely occupied that other molecules offer negligible interference with its thermal movement. The only obstacles to movement of the water molecule when *ϕ* → 0 comes from its interactions with the fixed molecular groupings comprising the walls of the channel. In a one-dimensional random walk, 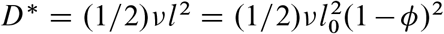, where *v* is the collision frequency, which in this model we assume independent of *ϕ*.

With these expressions for *K* and *D*^*\**^, we arrive at a formula for the diffusion permeability,

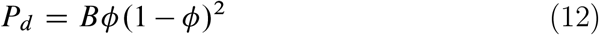

where for the prefactor,

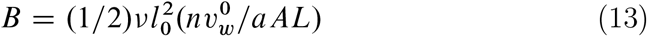

The dependence of *P*_*d*_ on *ϕ* is shown in Figure 2. When *ϕ* → 0, the permeability tends to zero. This behavior is reasonable, since water molecules cannot cross the membrane if the membrane holds no water. The permeability rises to a maximum value at *ϕ* = 1/3, then falls back to zero at *ϕ* = 1, where the hard spheres are touching and diffusional movement in the channel is suppressed. This condition is never achieved for temperatures above zero, and the mean field calculation given here should be understood as meaningful only if *ϕ* is not too close to unity.

**Figure 2.**
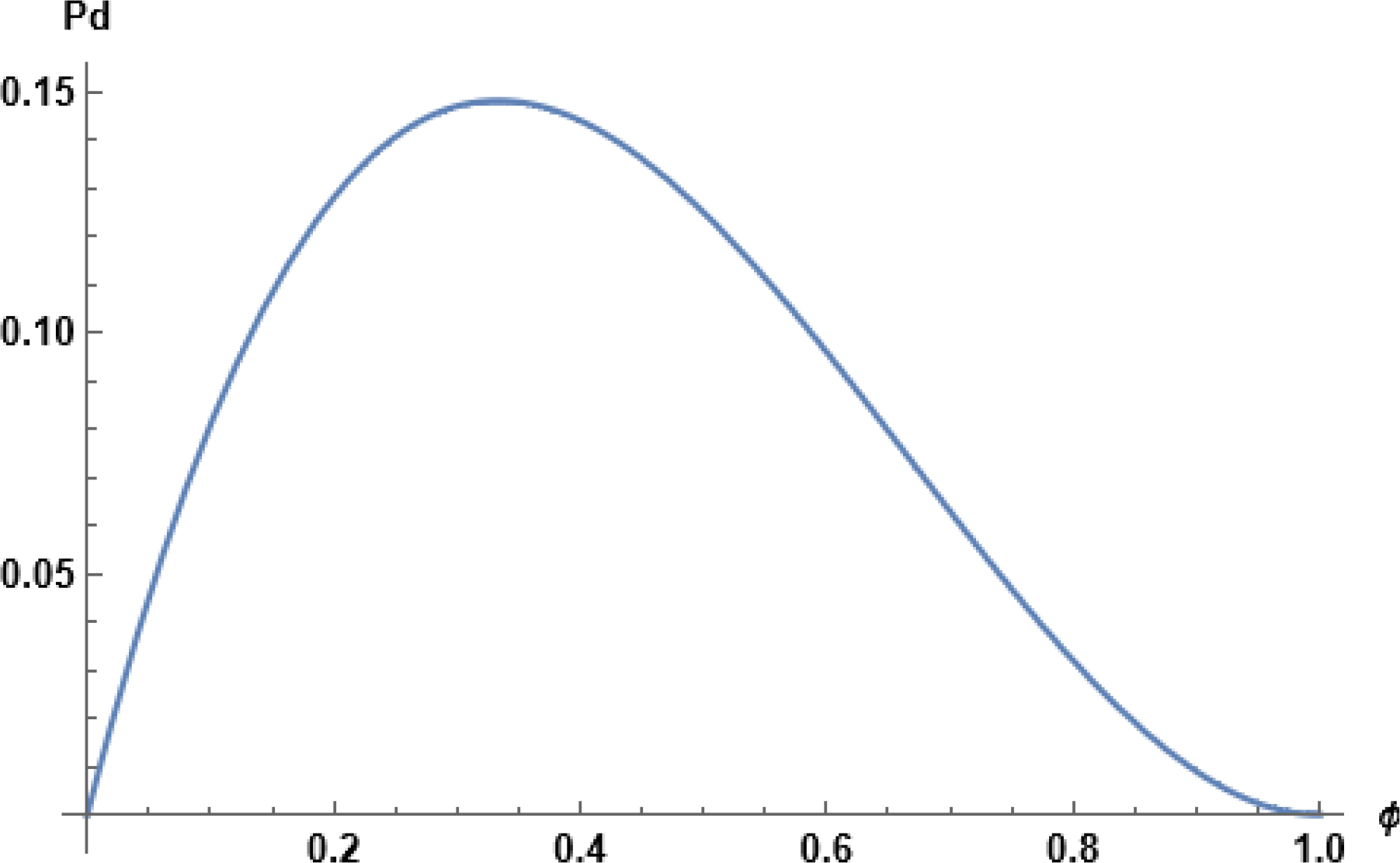
The diffusion permeability normalized to the prefactor B.

With *P*_*d*_ in hand and Eq. (11) for the *P*_*f*_ */P*_*d*_ ratio, the result for *P*_*f*_ follows,

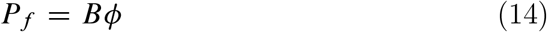

Thus *P*_*f*_ rises linearly with *ϕ* from zero at zero occupancy (for the same physical reason that *P*_*d*_ vanishes there), to its maximum value *B* at *ϕ* = 1.

Since *P*_*d*_ vanishes at *ϕ* = 1 in this model, an interesting interpretation of the course of the *P*_*f*_ */P*_*d*_ ratio as a function of *ϕ* (see Figure 1) is that it describes a transition from all-diffusive flow at low densities, *P*_*f*_ */P*_*d*_ → 1, to all-convective flow at *ϕ* = 1. The critical point for the transition is at *ϕ* = 1, where the diffusive component disappears. Of course, we know that one-dimensional systems do not exhibit phase transitions, which again shows the limitations of our mean-field interpretation of *ϕ*.

We can arrive at a numerical result if we convert *P*_*f*_ to the osmotic permeability per pore *p*_*f*_ D (*A/n*)*P*_*f*_ [1]. For the gramicidin A pore, *p*_*f*_ has been measured in two laboratories by distinct procedures as 1×10^−14^ cm^3^/s and 6×10^−14^ cm^3^/s. In our model, the combination 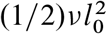 is the self-diffusion constant for water molecules in the pore under conditions of low fractional occupancy. If we use for it the value of the self-diffusion constant of bulk water, along with *ϕ* = 0:57 (see discussion involving gramicidin A in section 3), the diameter *a* of a water molecule, and *L*= 2.5 nm for the length of the gramicidin A pore, we get 5.6×10^−14^ cm^3^/s for *p*_*f*_ from Eq. (14).

## 5 Discussion

For water transport across lipid bilayers, the *P*_*f*_ */P*_*d*_ ratio is found to be equal to unity, as expected for diffusive movement of sparsely distributed water molecules dissolved in an oily environment [1,3]. For synthetic membranes more receptive to liquid water, the ratio is much greater than unity, indicating the dominance of bulk liquid flow [8, 9]. There was an early focus, currently all but lost, on the interesting question of why the *P*_*f*_ */P*_*d*_ ratio is greater than unity for water molecules moving in single file across narrow protein channels spanning biological membranes. The answer was that *P*_*f*_ */P*_*d*_ = *N*, the number of water molecules in the channel, as a seemingly reasonable condition if the passage of one molecule requires passage of all molecules ahead of it [1].

These early workers made no use of the parallel development in the theory of liquids of one-dimensional models that could be solved exactly [16–18]. In fact, the no-pass condition is inherent in these liquid models, as well as in the calculations of this paper. For example, Gürsey’s evaluation of the partition function integral for a one-dimensional fluid explicitly uses the condition that the coordinate *x*_*i*_ of the *i*th molecule is integrated only between the coordinates *x*_*i*-*1*_ and *x*_*i*C*1*_ of the two neighboring molecules [16]. Another example is the mean free path used in this paper, equal to zero at full occupancy.

Even though in developing our analysis we have made use of the no-pass condition, we did not arrive at the answer *N* for the single file *P*_*f*_ */P*_*d*_ ratio. Our wish for this paper is that it rekindle interest in a long-standing question remaining unresolved to this day [19].

## 6 Acknowledgments

The author is grateful to Alan Kay for many interesting discussions.

## 7 Data Availability Statement

No Data associated in the manuscript.

